# Estrogen attenuates the growth of human papillomavirus positive epithelial cells

**DOI:** 10.1101/2020.01.16.909986

**Authors:** Molly L. Bristol, Claire D. James, Xu Wang, Christian T. Fontan, Iain M. Morgan

**Affiliations:** Virginia Commonwealth University, Philips Institute for Oral Health Research, School of Dentistry; VCU Massey Cancer Center

## Abstract

Human papillomaviruses (HPVs) are small, double-stranded DNA viruses that are significant risk factors in the development of cancer, and HPV accounts for approximately 5% of all worldwide cancers. Recent studies using data from The Cancer Genome Atlas (TCGA) have demonstrated that elevated levels of estrogen receptor alpha (ERα) are associated with improved survival in oropharyngeal cancers, and these elevated receptor levels were linked with human papillomavirus positive cancers (HPV+cancers). There has been a dramatic increase in HPV-related head and neck squamous cell carcinomas (HPV+HNSCCs) over the last two decades and therapeutic options for this ongoing health crisis are a priority; currently there are no anti-viral therapeutics available for combating HPV+cancers. During our own TGCA studies on head and neck cancer we had also discovered the overexpression of ERα in HPV+cancers. Here we demonstrate that 17β-estradiol (estrogen) attenuates the growth/cell viability of HPV+cancers *in vitro*, but not HPV negative cancer cells. In addition, N/Tert-1 cells (foreskin keratinocytes immortalized with hTERT) containing HPV16 have elevated levels of ERα and growth sensitivity following estrogen treatment when compared with parental N/Tert-1. Finally, we demonstrate that there are potentially two mechanisms contributing to the attenuation of HPV+ cell growth following estrogen treatment. First, estrogen represses the viral transcriptional long control region (LCR) downregulating early gene expression, including E6/E7. Second, expression of E6 and E7 by themselves sensitizes cells to estrogen. Overall our results support the recent proposal that estrogen could be exploited therapeutically for the treatment of HPV positive oral cancers.

**Importance:** Human papillomaviruses cause around 5% of all human cancers, yet there are no specific anti-viral therapeutic approaches available for combating these cancers. These cancers are currently treated with standard chemo-radiation therapy (CRT). Specific anti-viral reagents are desperately required, particularly for HPV+HNSCC whose incidence is increasing and for which there are no diagnostic tools available for combating this disease. Using data from The Cancer Genome Atlas (TCGA) ourselves and others determined that the estrogen receptor α (ERα) is overexpressed in HPV+HNSCC, and that elevated levels are associated with an improved disease outcome. This has led to the proposal that estrogen treatment could be a novel therapeutic approach for combating HPV+cancers. Here we demonstrate that estrogen attenuates the growth of HPV+epithelial cells using multiple mechanisms, supporting the idea that estrogen has potential as a therapeutic agent for the treatment of HPV+HNSCC.

## Introduction

HPV is the most common sexually transmitted infection in the United States, infecting nearly every sexually active person at some point in their lives(1–8). Of the high-risk HPVs known to cause cancers, HPV16 is the most common genotype, accounting for 50% of cervical cancers and nearly 90% of HPV+HNSCCs(4, 9, 10). The level of HPV-related HNSCCs has become an epidemic in the last decade, with over half a million new cases per year worldwide(11). While prophylactic vaccines should be successful in preventing future HPV infections, there are currently no HPV-specific antiviral drugs to treat current HPV infections or HPV+HNSCC.

A number of studies have implicated steroid hormones, including 17β-estradiol (estrogen), as co-factors in HPV carcinogenesis(12–17). For example, the estrogen receptor has been shown to play an important role in the development of cervical cancer in a K14-HPV16 E7 transgenic mouse model, where estrogen was determined to work as a co-carcinogen with E7(14–16, 18, 19). However, the role of estrogen in the development of head and neck cancer in these transgenic mouse models has not been reported. In contrast to these results, studies demonstrate that high ERα expression correlates with increased survival in HPV+HNSCC(20, 21). These reports suggest ERα as a diagnostic marker but also raise the possibility of using estrogen as a therapeutic for the treatment of HPV+HNSCC. In support of the potential therapeutic potential of estrogen for HPV+ cancers, HeLa cells, an HPV18+ cervical cancer cell line, are extremely sensitive to estrogen treatment(22, 23). Given these recent reports we investigated the ability of estrogen to regulate the growth of HPV+ cell lines.

Analysis of our TCGA data agreed with those of others; the ERα receptor was overexpressed in HPV+HNSCC when compared with HPV-HNSCC, and higher expression predicted better overall survival(20, 21, 24). Here we report that estrogen treatment results in growth attenuation of HPV16+HNSCC lines (SCC47 and UMSCC104) but does not significantly alter the growth of HPV negative cancer cell lines. Previously we reported the transcriptional reprogramming of N/Tert-1 cells (foreskin cells immortalized by hTERT) by HPV16 (N/Tert-1+HPV16) and demonstrate here that the growth of these cells is attenuated by estrogen while control parental N/Tert-1 cell growth was not affected by estrogen treatment. We also treated human tonsil keratinocytes that were immortalized by HPV16 (HTK+HPV16) and these were severely growth attenuated following estrogen treatment. In SCC47, UMSCC104, UMSCC152, N/Tert-1+HPV16 (clonal and pooled lines), and HTK+HPV16 treated with estrogen, a significant reduction of early genes RNA transcript levels, including E6 and E7, is observed. Using HPV16-LCR (the long control region that regulates transcription from the HPV16 genome) luciferase vectors we demonstrate that estrogen can downregulate transcription from the HPV16 LCR. This down regulation has the potential to increase the p53 and pRb levels in the cells (the cellular targets for E6 and E7 respectively that promote degradation of these tumor suppressors). However, while p53 levels were altered in SCC47 and UMSCC104 cells, it was not altered in other lines; similarly, pRb was only significantly altered in HeLa cells, indicating the story may be more complex. While PARP1 cleavage was observed in SCC47, UMSCC152 and HeLa cells, it was not significantly altered in UMSCC104 cells, suggesting that growth attenuation is mediated by both apoptotic and non-apoptotic mechanisms, depending on the cell line. Finally, we treated N/Tert-1 cells expressing E6, E7 or E6+E7 (generated using retroviral transduction of the viral genes) with estrogen and demonstrate that expression of these viral oncoproteins by themselves results in growth attenuation of N/Tert-1 cells following estrogen treatment, however this growth attenuation is delayed when compared to N/Tert-1+HPV16 cells(25). Moreover, in these E6, E7, or E6+E7 cells the viral oncogene expression is not driven by the LCR and the levels of the viral RNA transcript do not change following estrogen treatment. In conclusion, the results demonstrate that estrogen attenuates the growth of HPV16+ keratinocytes and HPV+ cancer cells, and that there are potentially dual mechanisms for this attenuation; repression of viral transcription via targeting of the LCR, and cellular reprograming of the host by E6/E7 that promotes the estrogen sensitivity. Our results support the idea that estrogen can be used as a potential therapeutic for the treatment of HPV+HNSCC. In further support of this idea, we demonstrate that estrogen plus radiation treatment of the HPV+HNSCC line, SCC47 results in an additive attenuation of cell growth. No such affect was observed in the control HPV-HNSCC line, HN30.

## Results

### Estrogen attenuates the growth of HPV16 positive head and neck cancer cell lines

We have reported differential gene expression between HPV16+HNSCC and HPV-HNSCC using data from TCGA(24). We further analyzed this and observed that the ERα receptor expression was increased in HPV16+HNSCC versus HPV-HNSCC; as we were doing these studies two other reports were published demonstrating the increased expression of ERα in HPV+HNSCC(20, 21). Moreover, these studies demonstrated that increased levels of ERα predicted better survival suggesting that this receptor may be of diagnostic significance and that estrogen could be a novel therapeutic for targeting HPV+HNSCC(20, 21). We investigated the protein expression levels of ERα in HPV positive and negative cancer cells (Figure 1A). It is clear from this figure that any minor differences in protein expression of the ERα do not appear to be solely dependent on the HPV status of the cell line. Nevertheless, we proceeded to treat SCC47, UMSCC104 (HPV16+HNSCC integrated and episomal, respectively), C33a (HPV negative cervical cancer cell line) and HN30 (HPV-HNSCC) with estrogen and monitored cellular growth over a 6-day period (Figure 1B). There was a significant attenuation of the growth with SCC47 (i) and UMSCC104 (ii) following treatment with estrogen, but not with C33a (iii) or HN30 (iv). Likewise, the HPV18+ HeLa cervical cancer cells were also grown in the presence or absence of estrogen. Strikingly, all the HeLa cells appeared to be dead with the estrogen treatment at 72 hours (Figure 1Ci) when trying to observe HeLa cell growth in the presence or absence of estrogen, rendering cell growth observation impossible. To further analyze estrogen treatment in HeLa, cells were treated with varying doses of estrogen for 48-hours, and subjected to a cell viability assay by monitoring ATP release via Cell Titer-Glo; as observed in (Figure 1Cii), estrogen significantly reduced HeLa cell viability at all doses tested. The recently published data by Li et al also observed this phenomena, and indicates that estrogen may provide a unique approach to attenuate the growth or to kill HPV+ cells(23).

**Figure 1.**
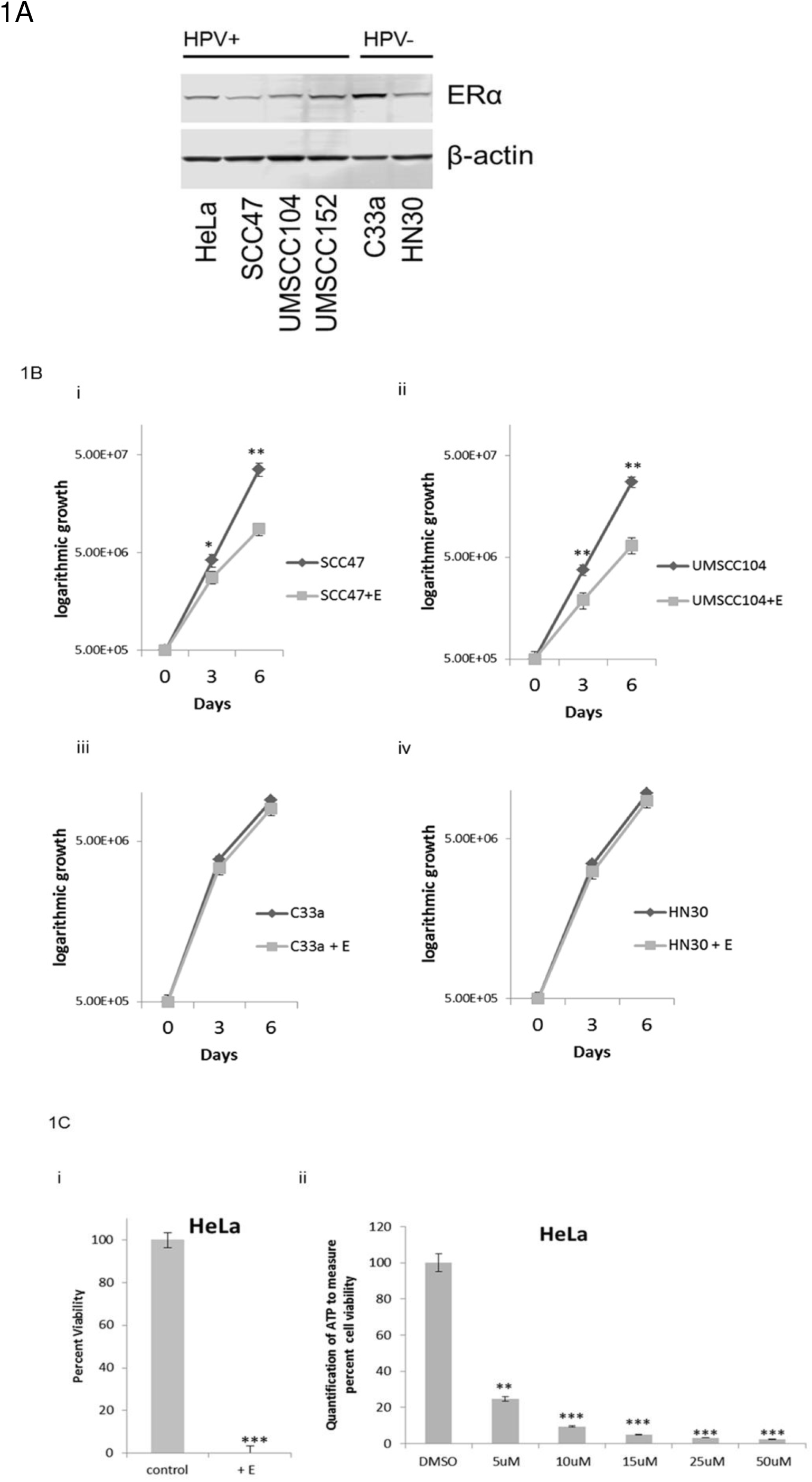
Estrogen attenuates the growth of HPV+ cancer cell lines. **A)** Cervical cancer cell lines HeLa and C33a, as well as HNSCC cell lines SCC47, UMSCC104, UMSCC152, and HN30 were analyzed for their expression of the ERα and compared to the loading control β-actin. HPV status is indicated above the blots. Experiments were conducted in triplicate and no significant correlation between HPV status and ERa expression was observed. **B)** HPV+ SCC47 **<u>(i)</u>** and UMSCC104 cells **<u>(ii)</u>**, and HPV-C33a **<u>(iii)</u>** and HN30 **<u>(iv)</u>** were seeded on day zero and grown in the presence or absence of 15µM estrogen. Cells were trypsinized and counted on day 3 and day 6 and cell counts are presented on a logarithmic scale. Statistical differences in both SCC47 and UMSCC104 cell can be observed at both day 3 and day 6. *p<0.05 **p<0.001. No statistical difference is observed between treatments on day 3 or day 6 in C33a **<u>(iii)</u>** or HN30 cells **<u>(iv)</u>**. Experiments were conducted in triplicate and error bars are representative of SE. **C) (i)** HeLa cells were grown in the presence or absence of 15µM estrogen for 72 hours then cells were counted for viability via trypan blue exclusion. **<u>(ii)</u>** Data is presented as % viability at 48 hours as measured by luciferase to monitor ATP via the Promega Cell Titer-Glo assay, over DMSO control. Experiments were conducted in triplicate and error bars are representative of SE. **p<0.001 **p<0.001.

We further investigated whether the estrogen treatment reduced the levels of HPV16 transcripts in these cells, as reduction of E6 and E7 levels have the potential to reactivate the p53 and pRb tumor suppressor pathways that would attenuate cellular growth. Figure 2A demonstrates that in SCC47, UMSCC104 and UMSCC152 (an HPV16+HNSCC line with a mixed population of integrated and episomal viral genomes) estrogen treatment for 7 days results in a significant reduction in viral RNA transcript levels. However, there was no significant reduction of the viral DNA levels in any of these cell lines during this treatment (Figure 2B). The results from Figures 1&2 demonstrate that estrogen can selectively attenuate the growth of HPV16+HNSCC cell lines and reduce the viral transcript levels in these cells.

**Figure 2:**
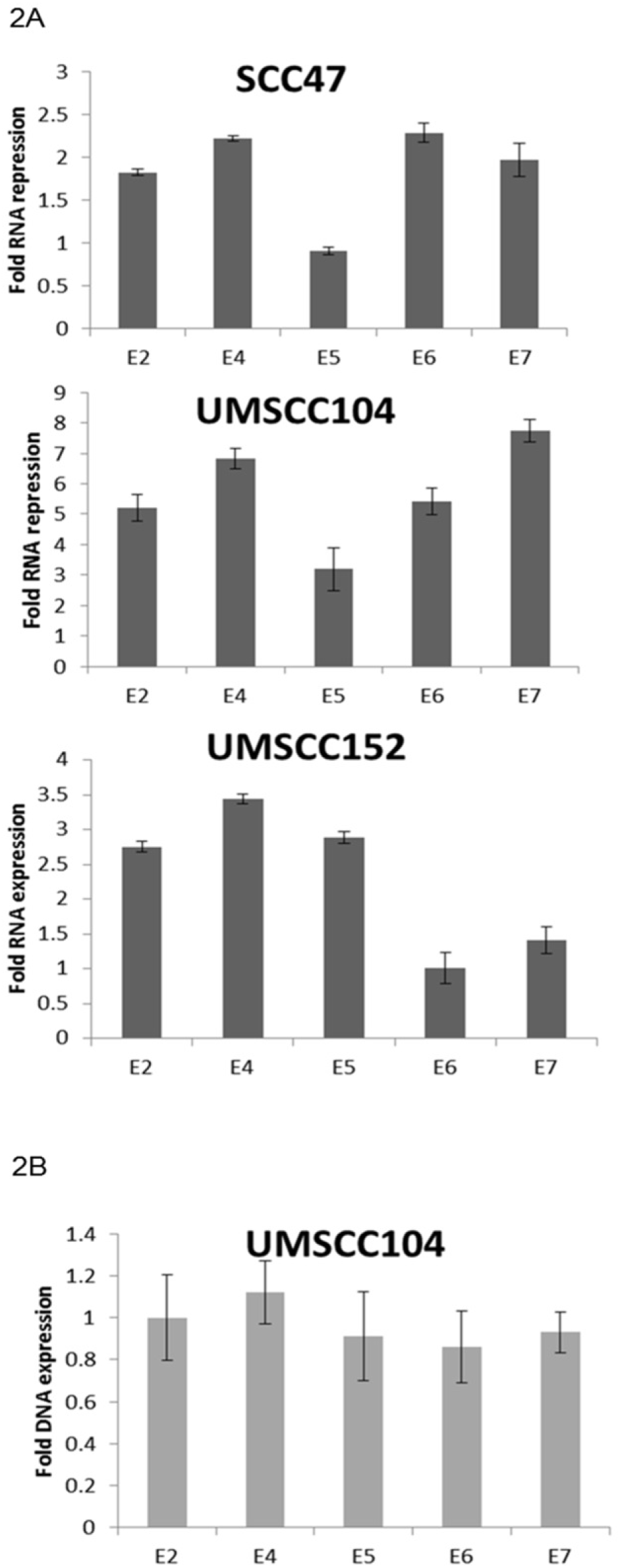
Estrogen significantly represses RNA expression of HPV16 early genes. **A)** SCC47, UMSCC104, and UMSCC152 cells were grown in the presence or absence of 15µM estrogen for 7 days. Cells were then harvested and RNA expression levels were monitored via qPCR for E2, E4, E5, E6 and E7, and compared to the loading control GAPDH. Data is presented as fold repression calculated from ΔΔCT calculated from the comparison of levels observed in control cells and further compared to GAPDH levels. **B)** Cells were treated as in A and DNA levels of E2, E4, E5, E6 and E7 levels were monitored via qPCR. Data is presented as fold repression calculated from ΔΔCT calculated from the comparison of levels observed in control cells and further compared to GAPDH levels. No significant DNA changes were observed in any of the cell lines and UMSCC104 is presented as representative data. Experiments were conducted in triplicate and error bars are representative of SE.

### An HPV16 isogenic model demonstrates that the presence of HPV16 imparts ERα upregulation and estrogen sensitivity

Previously we reported on the development of an HPV16 life cycle model in N/Tert-1 cells(24, 25). In N/Tert-1+HPV16 cells there is an increase in ERα expression over that in the parental N/Tert-1 cells (Figure 3A). The comparison between N/Tert-1 parent cells and N/Tert-1+HPV16 cells allows an isogenic comparison of their response to external reagents. Figure 3B demonstrates that control N/Tert-1 cell growth was not significantly affected by estrogen treatment over a 6-day period; in comparison, both pooled and clonally generated N/Tert-1+HPV16 cells were growth attenuated with estrogen treatment (Figure 3C). We also have investigated HPV16 host gene regulation in human tonsil keratinocytes immortalized by HPV16 (HTK+HPV16) and the growth of this cell line is severely attenuated by estrogen (Figure 3D)(26). Expression of the viral RNAs were downregulated by estrogen treatment in both N/Tert-1+HPV16 and HTK+HPV16 cells (Figure 3E). This is similar to the downregulation of viral RNA expression in the HPV16+HNSCC lines (Figure 2A).

**Figure 3:**
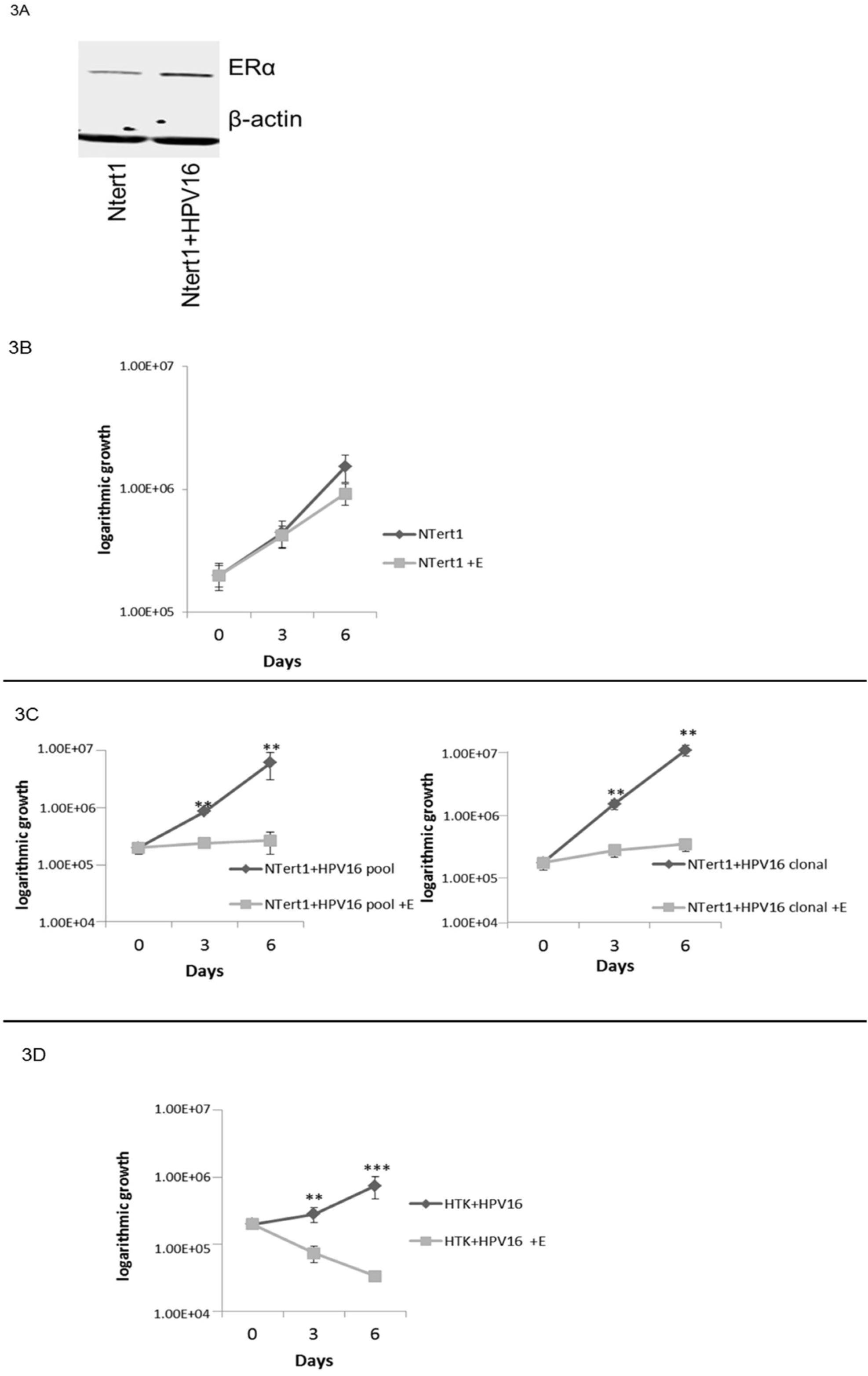
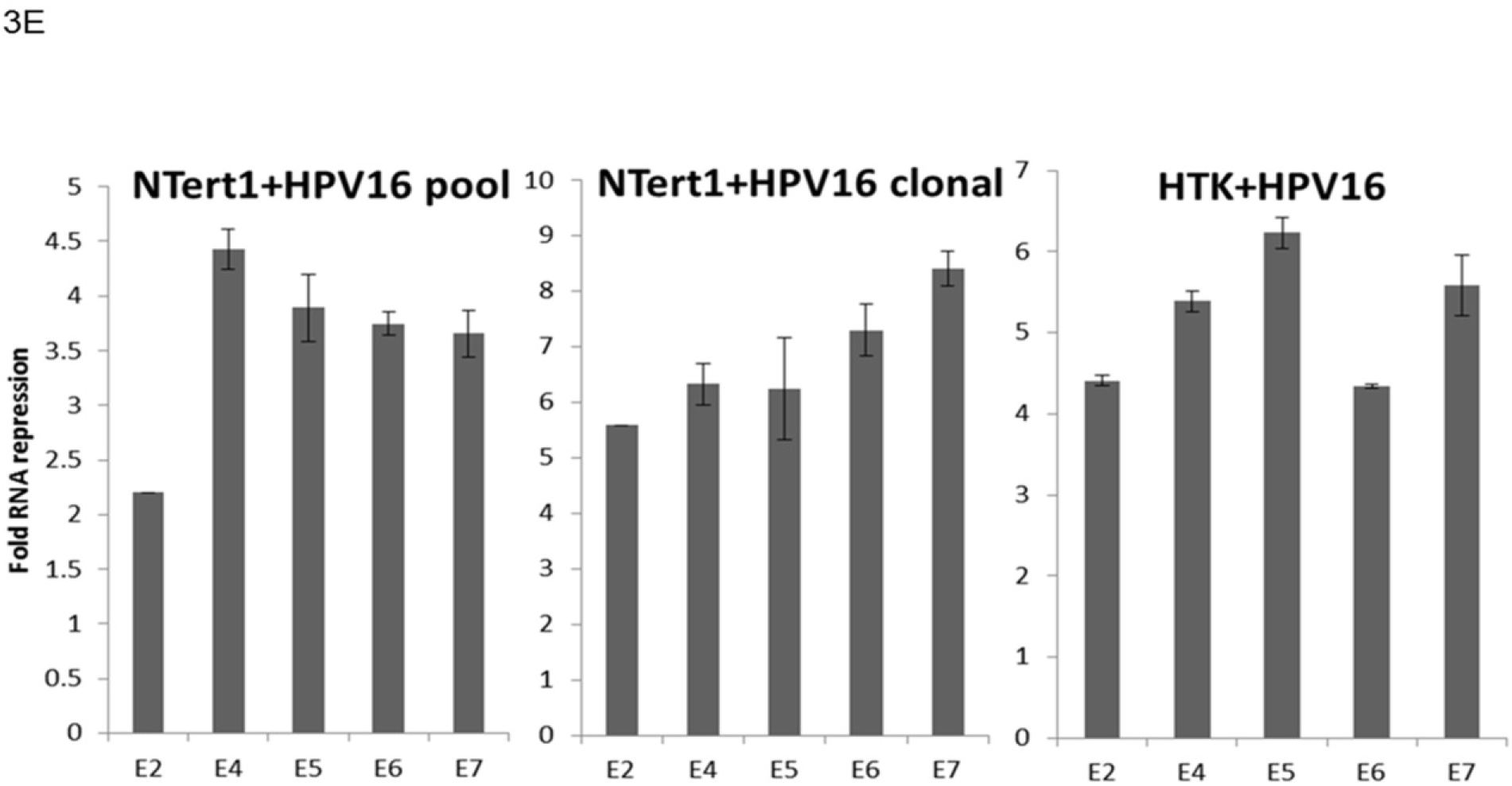
HPV16 confers estrogen sensitivity onto N/Tert-1 cells. **A)** Parental N/Tert-1 cell lines and our clonal N/Tert-1+HPV16 cells lines were analyzed for their overall ERα expression levels and compared to the loading control β-actin. **B-D)** N/Tert-<u>1 (**B**),</u> N/Tert-1+HPV16 (pool and clonal) (**C**), and HTK+HPV16 (**D**) cells were seeded on day zero and grown in the presence or absence of 15 trypsinized and counted on day 3 and day 6 and cell counts are presented on a logarithmic scale. Statistical differences can be observed at both day 3 and day 6 in all lines except the parental N/Tert-1 cells. **p<0.001 ***p,0.0001. Experiments were conducted in triplicate and error bars are representative of SE. **E)** Pooled N/Tert-1+HPV16, clonal N/Tert-1+HPV16, and pooled HTK+HPV16 cells were grown in the presence or absence of 15µM estrogen for 7 days. Cells were then harvested and RNA expression levels were monitored via qPCR for E2, E4, E5, E6 and E7, and compared to the loading control GAPDH. Data is presented as fold repression calculated from ΔΔCT calculated from the comparison of levels observed in control cells and further compared to GAPDH levels. Experiments were conducted in triplicate and error bars are representative of SE.

### Estrogen represses transcription from the HPV16 long control region (LCR)

Figures 2&3 demonstrate that estrogen treatment of HPV16+ cells results in the repression of viral RNA expression. Transcription of HPV16 viral genes is regulated by the Long Control Region (HPV16-LCR), a region that is regulated by a number of host transcription factors. We constructed a reporter plasmid where luciferase gene expression is regulated by the HPV16-LCR (pHPV16-LCR-Luc), transfected this vector into C33a cells, and monitored transcription levels of the pHPV16-LCR-Luc via relative fluorescence units (RFU) in the presence or absence of estrogen. Estrogen treatment resulted in a significant reduction of luciferase expression (Figure 4A), while expression from a control luciferase plasmid (pgl3 basic) was not affected by estrogen treatment. Because of the effects observed in HeLa cells (Figure 1C), we sought to determine if the LCR repression was also observed in HPV18 used a previously described pHPV18-LCR-luc plasmid(27); similar significant repression of the HPV18-LCR was also observed (Figure 4B). We carried out similar experiments in N/Tert-1 cells where estrogen treatment also significantly reduced luciferase activity in cells transfected with pHPV16-LCR-Luc (Figure 4C), but did not reduce the control luciferase plasmid. The conclusion from Figures 2–4 is that estrogen represses transcription from the HPV16 long control region to downregulate expression of early viral genes.

**Figure 4:**
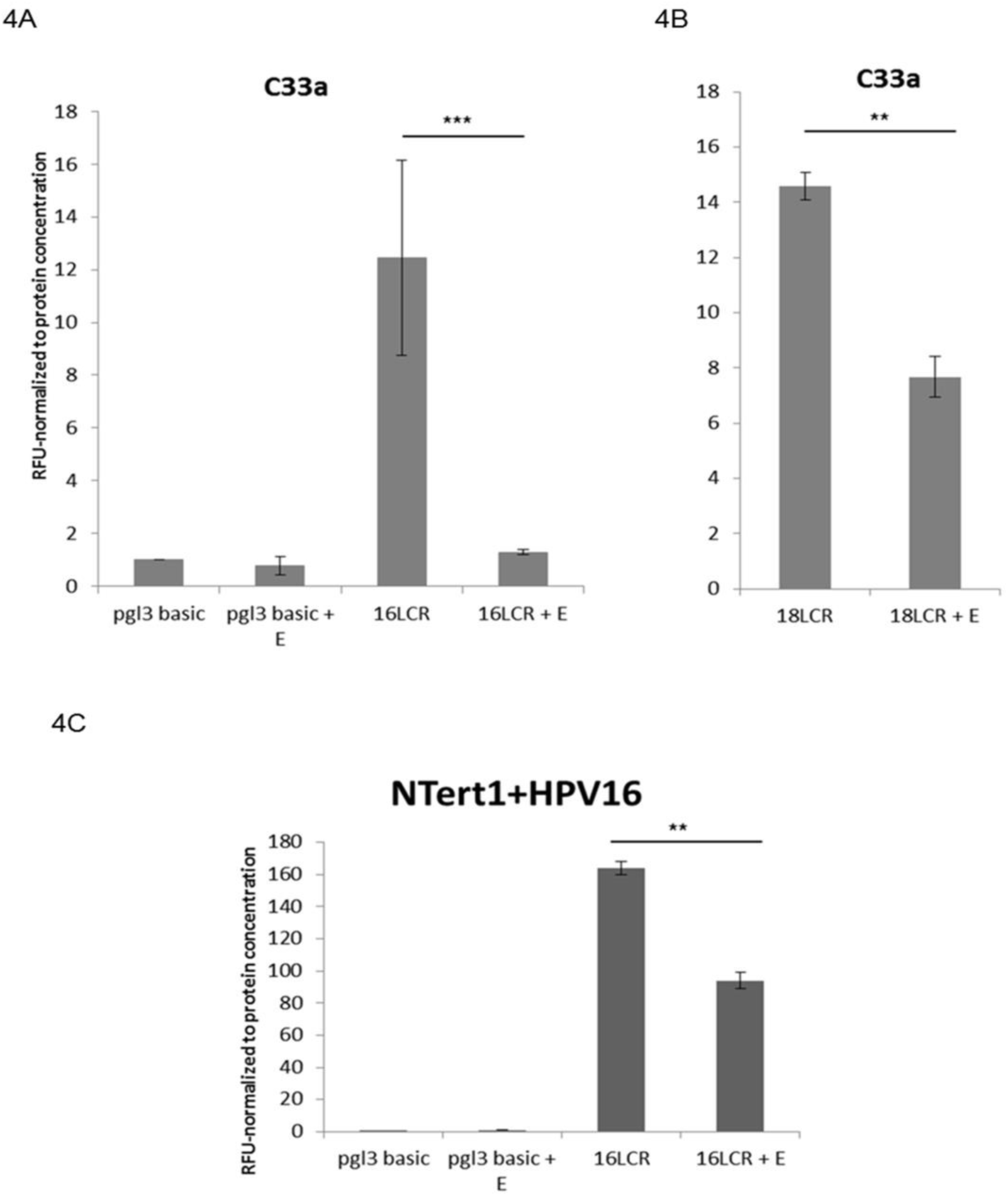
Estrogen significantly represses HPV16 and HPV18 LCR transcription. **A)** C33a cells were transfected with 1µg of pgl3 basic backbone (control), 1µg 16LCR-pGL3, or in **B)** 1µg 18-LCR-pGL3 and grown in the presence or absence of 15µM estrogen. **C)** N/Tert-1 cells were transfected with1µg of pgl3 basic backbone, or 1µg 16LCR-pGL3 and grown in the presence or absence of 15µM estrogen. Forty-eight hours post transfection, a luciferase-based assay was utilized to monitor levels of LCR transcription. Data is presented as RFU, normalized to total protein concentration as monitored by a standard BSA assay. **p<0.001 ***p<0.0001

### Estrogen increases DNA damage and initiates apoptosis in some HPV+ cancer cells

Downregulation of E6 and E7 expression by estrogen could result in the elevation of p53 and pRb expression (their respective tumor suppressor targets)(28–41). Previously, studies have shown that when E2 is overexpressed in HPV positive cervical cancer cells it represses transcription from the viral LCR and this repression reduces E6 and E7 levels and reactivates the p53 and pRb tumor suppressor proteins(31, 42–44, 44–50). Moreover, E2 overexpression and loss of E6/E7, results in the elevation of p53 and pRb that allows for previously observed attenuation of growth in HeLa cells(23, 31, 45, 46, 48). Similarly, our studies indicate estrogen treatment represses transcription from the LCR to reduce expression of E6 and E7 levels. We therefore analyzed the protein levels of p53 and pRb in our cancer cell lines in the presence or absence of estrogen, as well as monitor γh2AX as a marker for the initiation of the DNA damage response(51), and the ratio of cleaved-PARP1/PARP1 as a marker for apoptosis. These western blots are presented in Figure 5A with accompanying densitometry analysis (Figure 5B).

**Figure 5:**
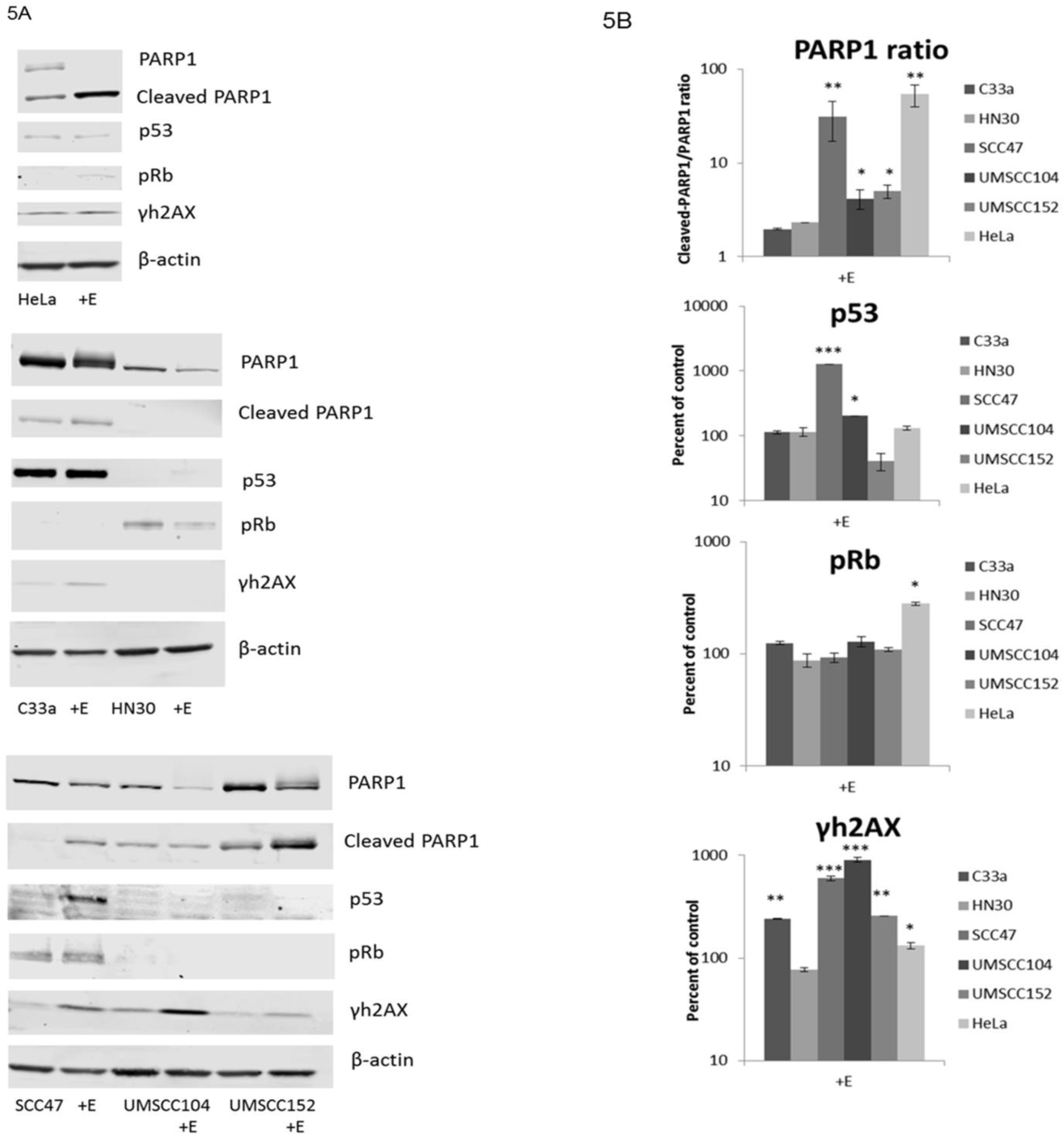
Estrogen alteration of protein expression in cancer cell lines. **A)** HPV18+ HeLa (top panel), HPV-C33a and HN30 (middle panel), and HPV16+ SCC47, UMSCC104, and UMSCC152 cells (bottom panel) were grown in the presence or absence of 15µM estrogen for 48 hours. Cells were then lysed and analyzed via western blot for PARP1, cleaved-PARP1, p53, pRb, and γh2AX. β-actin was used as a loading control. **B)** Densitometry analysis was compared from three independent experiments. For PARP1 the ratio of cleaved to non-cleaved PARP1 is given and the rest are presented in graphs as percent of control cells. All are normalized to loading control, and are in log scale.

As expected, analysis of the response to estrogen in the sensitive HeLa cells revealed a significant increase in p53, pRb, γh2AX, and PARP1 cleavage (Figure 5A, top panel). Confirming the previously observed increase in apoptosis following estrogen in HeLa cells(23). Furthermore, analysis in the HPV-cancer cells reveals no dramatic alterations in p53, pRb, or PARP1 cleavage; however, there is a significant increase in γh2AX in C33a cells (Figure 5A, middle panel). This increase in γh2AX reveals that estrogen is still initiating DNA damage; however, it appears that this damage alone is not sufficient to inhibit the growth of the C33a cells. Western blot analysis of our HPV+HNSCC lines reveals a less than clear cut mechanism that allows for the reduction in cell growth observed (Figure 5A, bottom panel). While all cells exhibited an increase in γh2AX and PARP1 cleavage indicating estrogen induces DNA damage that results in an increase of apoptosis, no significant alterations in pRb were observed in any of our HPV+HNSCC lines, and p53 was only significantly increased in SCC47 and UMSCC104 cells. Therefore, the reactivation of these tumor suppressors following estrogen treatment does not fully explain the attenuation of cell growth in the HPV16+ cells.

### Expression of the viral oncogenes promotes delayed cell growth attenuation following estrogen treatment

We next investigated whether the transcriptional reprograming of N/Tert-1 cells carried out by HPV16 oncogenes alone could sensitize cells to estrogen and attenuate cellular growth. To do this we expressed E6 or E7 or E6+E7 in N/Tert-1 cells and further compared these cells to those expressing the full HPV16 genome (N/Tert-1+HPV16); these E6, E7, and E6+E7 cell lines were generated using retroviral delivery and have been described previously(26, 52). Figure 6A demonstrates again that in N/Tert-1 control cells, estrogen treatment does not attenuate cellular growth (Figure 6Ai) but the presence of the entire HPV16 genome promotes such attenuation (Figure 6Aii). The presence of E6, E7 or E6+E7 resulted in growth attenuation following estrogen treatment (Figures 6Aiii-v), although it was not observed on day 3, instead delaying the attenuation of cell growth that was observed with the entire HPV16 genome (comparison of Day 3 is normalized and presented in Figure 6B). As the expression of the E6 and E7 in panels 5Aiii-v is not driven from the viral LCR, but rather from retroviral sequences, we anticipated that the RNA levels of the oncogenes would not be regulated by estrogen. This is indeed the case; estrogen treatment did not alter E6 or E7 levels in the cells transduced with the retroviral vectors (Figure 6C). Therefore, the growth attenuation of these cells following treatment with estrogen can be contributed to the expression of the viral oncoproteins, and likely due to the transcriptional reprograming of these cells carried out by these proteins.

**Figure 6:**
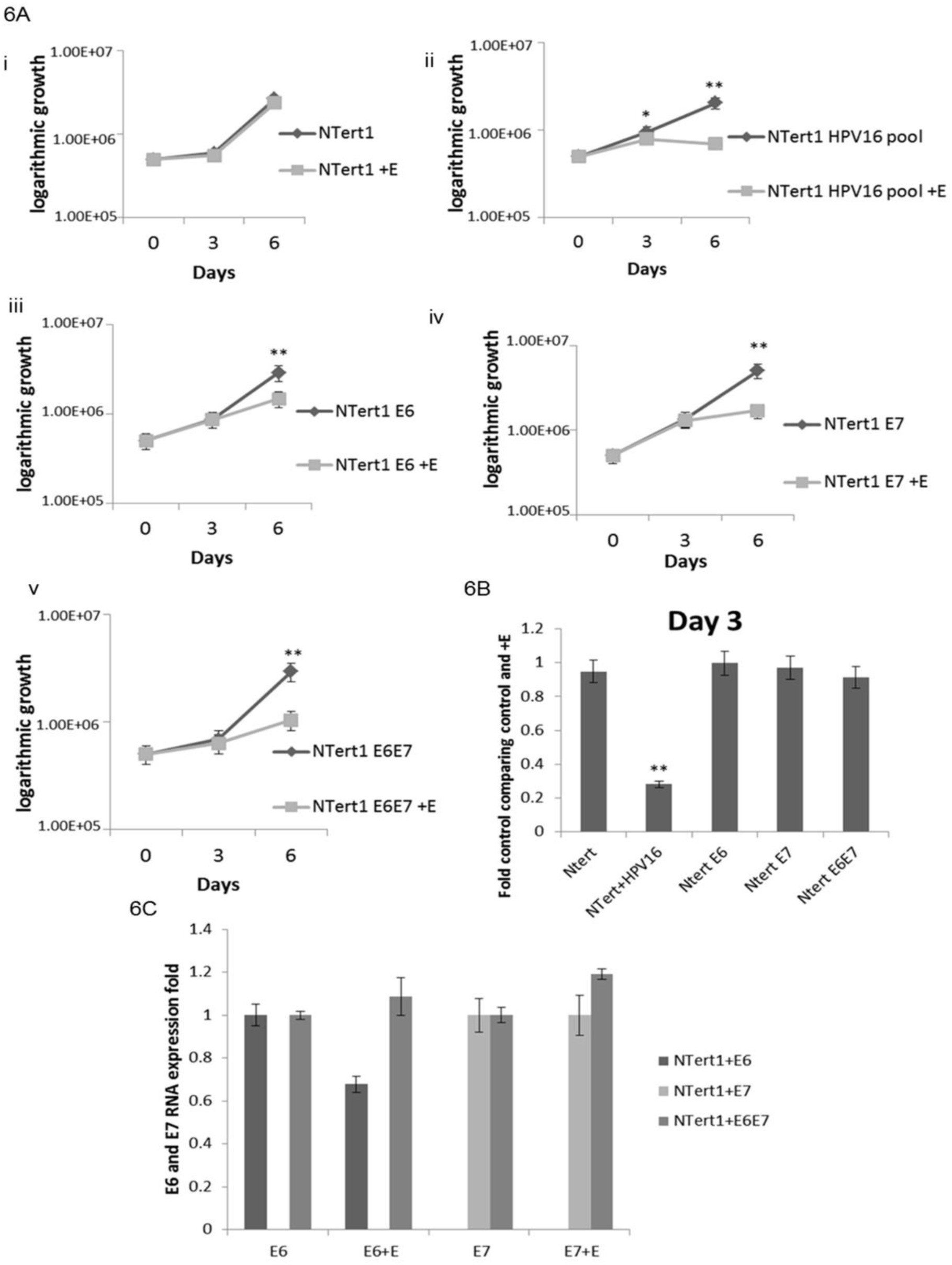
E6 and E7 expression by themselves sensitizes N/Tert-1 cells to estrogen. **A (i)** N/Tert-1, **(ii)** N/Tert-1+HPV16, **(iii)** N/Tert-1+E6, **(iv)** N/Tert-1+E7, and **(v)** N/Tert-1+E6E7 cells were seeded on day zero and grown in the presence or absence of estrogen. Cells were trypsinized and counted on day 3 and day 6 and cell counts are presented on a logarithmic scale. Statistical differences can be observed at both day 3 and day 6 **(ii)**, but only on day 6 in **(iii-v)** *p<0.05 **p<0.001. **B)**. Day 3 cell counts are compared as percent of control and normalized, only N/Tert-1+HPV16 presents statistical difference at this time point. **p<0.001. **C)** N/Tert-1+E6, N/Tert-1+E7, and N/Tert-1+E6E7 cells were analyzed for their RNA expression levels of E6 and E7, and compared to the loading control GAPDH. Data is presented as fold expression as calculated from ΔΔCT calculated from the comparison of levels observed in control cells and further compared to GAPDH levels. No statistical differences were found.

### Estrogen and radiation treatment of HPV positive and negative cancer cells

Radiation treatment is a standard of care therapy for HPV+HNSCCs. We treated C33a, HN30 and SCC47 cells with estrogen and then treated them with 2, 5 and 10 Gy of radiation to investigate whether estrogen can promote further response to this treatment. For C33a (Figure 7A) and HN30 (Figure 7B), the presence of estrogen made no significant difference to the response to radiation treatment. For SCC47 cells, treatment with estrogen by itself attenuated cell growth, as shown in Figure 1Bi. As observed in Figure 7C, treatment with radiation did not have a dramatic effect on the growth of SCC47. However, because SCC47 cells were highly sensitive to estrogen alone, the additive effect observed with estrogen and radiation lead to ∼80% loss in cell viability even at 2 Gy radiation. This is promising and suggests that estrogen treatment may provide a unique opportunity to allow for increased responsiveness to radiation treatment in the clinic at reduced radiation doses for HPV+HNSCC.

**Figure 7:**
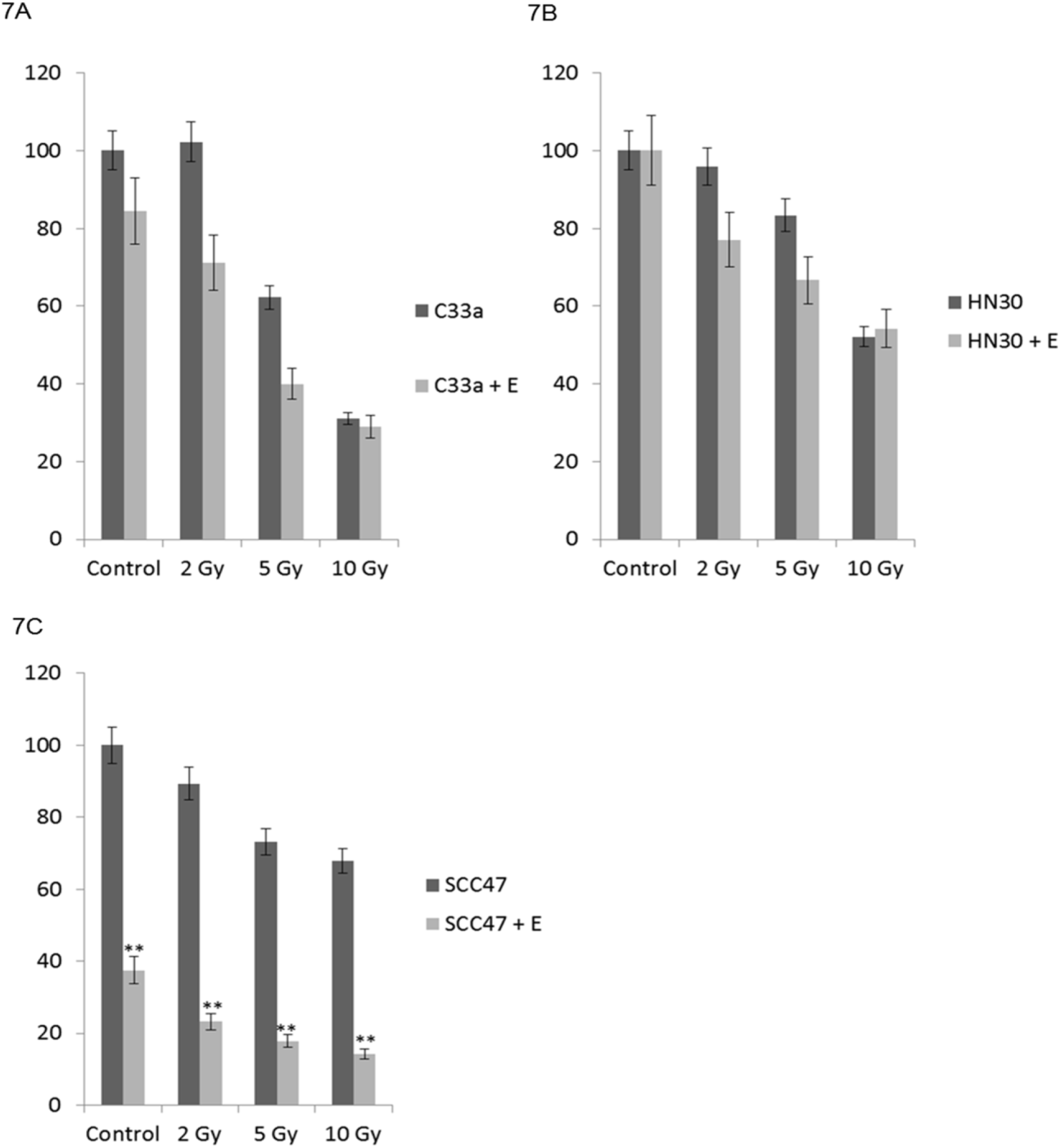
Estrogen enhances the response to radiation in SCC47 cells but not in C33a or HN30 cells. **A)** C33a **B)** HN30, and **C)** SCC47 cells were maintained in estrogen for 72 hours. Noted cells were then radiated with 2, 5, or 10 Gy radiation, and cells were trypsinized and counted by trypan blue exclusion for viability 72-hours post-irradiation. Data is presented as % viability compared with untreated control cells. Experiments were conducted in triplicate and error bars are representative of SE. **p<0.001.

## Discussion

While the prophylactic vaccine should decrease the incidence of HPV in the upcoming decades, we currently lack antiviral treatments to target those already infected with the virus. Likewise, HPV-related HNSCC are on the rise and this oncogenic virus has bypassed tobacco as the main carcinogen in the oropharyngeal region(3, 11, 53). Despite HPV+ and HPV-HNSCCs being very different both phenotypically and genotypically in terms of their pathological and molecular mechanisms of carcinogenesis and in their response to therapy, they are still treated the same in the clinic(54). It is therefore of particular interest to develop HPV-specific treatments for HPV+HNSCC.

Analysis of TCGA data showed that the expression of the estrogen receptor alpha (ERα) was highly significantly upregulated in HPV16+HNSCC vs HPV-HNSCC(20, 21, 24). The ERα also decreased as stages advance, so we initially rationalized that estrogen may play a role in the early development of cancer. This differential expression of the ERα presented an opportunity to exploit a significant difference between HPV+/-HNSCC and to possibly develop a specific targeted approach. Our initial hypothesis aligned with previous indications that the estrogen and the ERα increase the risk of cervical cancer, and we further predicted that high doses of estrogen would initiate the DNA damage response (DDR) (14–16, 18, 19, 55–58). Based on our previously published data, we further predicated that this increase in the DDR via estrogen would enhance HPV tumorigenicity and ultimately result in worse outcomes and disease progression(59). However, it soon became clear that our initial hypothesis was incorrect when HPV+ cells were specifically sensitized via estrogen treatment, while HPV-cells showed little to no response. We were also extremely surprised with the dramatic response to estrogen that we observed in HeLa cells, although recently published data now confirms our observations(23). This recent study utilizing HeLa cells as a model to analyze steroid signaling confirmed these cells are particularly sensitive to estrogen. Li et al showed that estrogen induced classical caspase-3-mediated apoptosis via a multi-step molecular mechanism, however this study did not take into account the HPV status of their cell model and may have missed an underlying viral mechanism by which estrogen was able to induce the cell death they observed(23). More specifically, HeLa cells are intrinsically dependent on the expression of E6 and E7(60); if estrogen is able to reduce viral levels of these vital oncoproteins, this could contribute to the rapid death progression observed in HeLa cells, although it likely not the only mechanism.

While the expression of the ERα was found to be upregulated in HPV+HNSCC, and via HPV expression in our N/Tert-1 model, we do not believe that the overall ERα expression level is the only reason that HPV+ cells are sensitive to estrogen. Among the cell lines we analyzed for estrogen sensitivity, the C33a cells had the highest protein level as observed by western blot (Figure 1A), yet C33a cells showed little to no cell growth response to estrogen alone (Figure 1Biii), however estrogen did increase γh2AX demonstrating these cells are responsive to estrogen (Figure 5A, middle panel), while only providing moderate sensitization to irradiation (Figure 7A). It is likely that estrogen/HPV specific interactions, both via the LCR and E6/E7, are responsible for the growth inhibition and cell death we observed in our HPV+ cell lines, not from DNA damage signaling alone. Nevertheless, the HPV upregulation of ERα likely ensures the ability of HPV infected cells to respond to estrogen treatment. Further expanding this, high expression of the ERα alone, as observed in C33a cells, is not enough to confer estrogen sensitivity; HPV upregulation of the ERα in conjunction with HPV specific estrogenic signaling, initiates a complex signaling cascade to initiate estrogen sensitivity.

HPV+HNSCC is most commonly associated with males, found at a 4:1 higher ratio than observed in females(61). While estrogen is typically associated with females, men do in fact express appreciable levels of the estrogen receptors and circulating estradiol levels in males are the same or higher than observed in post-menopausal women(62–65). Therefore, this could begin to explain some of the sex related differences observed in the instances of HPV+HNSCC and presents an interesting observation for future studies.

It isn’t clear what control region in the HPV16 LCR is responsible for transcriptional repression following estrogen treatment. However, it has been shown that the ERα can interact with AP1 via c-Jun and there are known AP1 binding sites in the HPV16 LCR that may mediate the response of this region to estrogen(66–72). This will be investigated in future studies.

Future studies determining the exact mechanism of the interaction between estrogen and HPV may provide additional opportunities to provide more specific targeted approaches to exploit this HPV specific sensitization to estrogen for therapeutic gain in the treatment of HPV+cancers. Overall our results indicate that estrogen may provide an approach that could be exploited therapeutically for the treatment of HPV+ epithelial cells.

## Materials and Methods

### Cell culture

C33a (ATCC), HN30 (generous gift from Dr. Hisashi Harada, VCU Philips Institute), SCC47 (Millipore), and HeLa (generous gift from Dr. Alison McBride, NIAID) cells were grown in Dulbecco’s modified Eagle’s medium (Invitrogen) and supplemented with 10% charcoal stripped fetal bovine serum (Gemini Bio-products). UMSCC104 (Millipore), and UMSCC152 (ATCC) cells were grown in Eagle’s Minimum Essential Medium (EMEM, Invitrogen) supplemented with non-essential amino acids (NEAA, Gibco) and 10% charcoal stripped fetal bovine serum. N/Tert-1 cells and all derived cell lines, as well as HTK+HPV16 cells (a generous gift from Dr. Craig Meyers, UPenn, Hershey) have been describe previously(24, 25, 52, 59) and were maintained in keratinocyte-serum free medium (K-SFM, Invitrogen), supplemented with a 1% (vol/vol) penicillin-streptomycin mixture (ThermoFisher Scientific). All N/Tert-1 cells were also supplemented with 4 μg/ml hygromycin B (Millipore Sigma). All cells not directly purchased from providers were cell type confirmed by Johns Hopkins or MD Anderson cell line authentication services, were maintained at 37°C in a 5% CO_2_–95% air atmosphere, routinely passaged every 3-4 days and routinely monitored for mycoplasma.

### Trypan blue exclusion

Cell supernatant was collected to allow for dead cell collection; attached cells were harvested by trypsinization and added to cell supernatant. Total cells were stained with trypan blue and viable cells counted. Total number of cells was recorded and viable cell ratio was calculated.

### Plasmids

pHPV16-LCR-Luc was generated by PCR amplification of the HPV16 LCR from W12 cells, introducing *Kpn*I and *Bgl*III restriction sites, and cloned into a pGL3 backbone (cloning primers listed below). The other plasmids utilized in these studies have been previously reported by others or used and described by this laboratory: PGL3 basic(73), pHPV18-LCR-Luc(27), HPV16 E6 (p6661 MSCV-IP N-HA only 16E6 – Addgene plasmid # 42603 Dr. Peter Howley), HPV16 E7 (p6640 MSCV-P C-FlagHA 16E7-Kozak - Addgene plasmid # 35018 – Dr. Peter Howley). HPV16 E6E7 (pLXSNE6E7 Addgene#52394 – Dr. Denise Galloway)

### pHPV16-LCR-Luc Cloning primers (Invitrogen)

HPV16 LCR-forward 1 (position 7153) 5’-TCGAG<u>GTACC</u>GCTGTAAGTATTGTATGT-3’; forward 2 (position 7288) 5’-TCGAG<u>GTACC</u>ATGCTTGTGTAACTATTG-3’; forward 3 (position 7423) 5’-TCGAG<u>GTACC</u>GTAGCGCCAGCGGCCATT-3’; forward 4 (position 7531) 5’-TCGAG<u>GTACC</u>GTACGTTTCCTGCTTGCC-3’; forward 5 (position 7668) 5’-TCGAG<u>GTACC</u>CACTATGCGCCAACGCCT-3’; forward 6 (position 7737) 5’-TCGAG<u>GTACC</u>GCATATTTGGCATAAGGT-3’; forward 7 (position 7873) 5’-TCGAG<u>GTACC</u>CACATTTACAAGCAACTT-3’; reverse (position 94) 5’-TCGA<u>AGATCT</u>GGGTCCTGAAACACTGCAGTTCTT-3’.

### Transfection Assays and Transcriptional Activity

Note cells were plated at 5 x 10^5^ in 100-mm dishes. The following day, plasmid DNA was transfected using the calcium phosphate method for C33a. N/Tert-1 cells were transfected utilizing lipofectamine 2000 (according to manufacturer’s instructions, ThermoFisher Scientific). 24-hours post transfection cells were washed and noted cells were supplemented with 15µM 17β-estradiol. 48-hours post transfection, cells were harvested utilizing Promega Reporter Lysis Buffer and analyzed for luciferase using the Promega Luciferase Assay System. Concentrations were normalized to protein levels, as measured by the BioRad Protein Assay Dye, and relative fluorescence units were measured using the BioTek Synergy H1 Hybrid Reader. Experiments were performed in triplicate.

### Western blots

Cells were trypsinized, washed twice with phosphate buffered saline (PBS), pelleted, then re-suspended in 200 μl of lysis buffer (0.5% Nonidet P-40, 50 mM Tris, pH 7.8, 150 mM NaCl) supplemented with a protease inhibitor mixture (Roche Molecular Biochemicals). The cell and lysis buffer mixture was incubated on ice for 30 min, centrifuged for 10 min at 18,000g at 4 °C, and supernatant was collected. Protein levels were determined utilizing the Bio-rad protein assay (Bio-rad). Equal amounts of protein were boiled in 4x Laemmli sample buffer (Bio-rad). Samples were then loaded onto a 4– 12% gradient gel (Invitrogen), ran at 120 V for ∼2 h and transferred at 100 V for 1 h onto nitrocellulose membranes (Bio-rad) using the wet blot method. The membrane was then blocked in Odyssey blocking buffer (diluted 1:1 with PBS), at room temperature for 1 h. After blocking, the membrane was probed with noted antibodies diluted in blocking buffer, and incubated O/N at 4 °C: p-histone H2A.X Rabbit 1:1000 (Cell Signaling #9718S), β-actin Mouse 1:2000 (Santa Cruz sc-81178), ERα Rabbit 1:1000 (AbCam ab32063), p53 Mouse 1:1000 (Cell Signaling 2524S), pRb Mouse 1:1000 (Cell Signaling 9309S), PARP1 Mouse 1:1000 (SantaCruz sc-8007), cleaved-PARP1 Rabbit 1:1000 (Cell Signaling 9541S). Following incubation with primary antibody, the membrane was washed with 0.01% PBS-Tween wash buffer before probing with Odyssey secondary antibody diluted 1:20,000, Goat anti-mouse IRdye 800 cw, Goat anti-rabbit IRdye680cw for one hour at room temperature. The membrane was then washed in 0.01% PBS-tween before infrared scanning using the Odyssey Li-Cor imaging system, also used to perform densitometry analysis. Experiments were performed in triplicate.

### SYBR green qRT-PCR

At the time of harvest, cells were washed twice with phosphate buffered saline (PBS). RNA was immediately isolated using the SV Total RNA Isolation System (Promega) following the manufacturer’s instructions. Two micrograms of RNA were reverse transcribed into cDNA using the High Capacity Reverse Transcription Kit (Applied Biosystems). cDNA and relevant primers were added to PowerUp SYBR Green Master Mix (Applied Biosystems) and real-time PCR performed using 7500 Fast Real-Time PCR System (Applied Biosystems). Results shown are the average of three independent experiments with relative quantity of genes determined by the ΔΔCt method normalized to the endogenous control gene GAPDH.

### *Primers* (Invitrogen)

GAPDH 5’-GGAGCGAGATCCCTCCAAAAT-3’ (forward) and 5’-GGCTGTTGTCATACTTCTCATGG-3’. E2 5’-TGGAAGTGCAGTTTGATGGA-3’ (forward) and 5’-CCGCATGAACTTCCCATACT-3’ (reverse). E4 5’-GGCACCGAAGAAACACAGAC-3’ (forward) and 5’-AATCCGTCCTTTGTGTGAGC-3’ (reverse). E5 5’-CACAACATTACTGGCGTGCT-3’ (forward) and 5’-ACCTAAACGCAGAGGCTGCT-3’ (reverse). E6 5’-AATGTTTCAGGACCCACAGG-3’ (forward) and 5’-GCATAAATCCCGAAAAGCAA-3’ (reverse). E7 5’-CCGGACAGAGCCCATTACAAT-3’ (forward) and 5’-ACGTGTGTGCTTTGTACGCAC-3’ (reverse).

### CellTiter-Glo protocol for measuring cellular ATP

2000 cells were plated in 200uL media in clear bottom black 96-well plates (Greiner Bio One, 655090). The following day, media was removed from cells and replaced with 200uL media containing 17β-Estradiol at differing concentrations. Cells were then incubated for 48-hours. Afterwards, 25uL of reconstituted CellTiter-Glo Luminescent Cell Viability reagent was added to each well and incubated for 5 minutes (Promega, G7571). Luminescence readings were taken using the BioTek Synergy H1 Hybrid Reader. Viability percentages were calculated by normalizing to DMSO treated cell readings utilizing media only wells for blanking. DMSO wells were normalized to 100%.

### Radiation

Noted cells were exposed to γ-IR using a ^137^Cs irradiator. Radiation treatment consisted of a single dose of irradiation at 2, 5, or 10 Gy. In our studies, cells were exposed to estrogen for 72 hours before irradiation. Post-irradiation, cells were washed once with PBS and medium replaced. Estrogen was then maintained on noted cells for an additional 72 hours before cells were trypsinized and counted for cell viability.

## Acknowledgments

This work was supported by the VCU Philips Institute for Oral Health Research and the National Cancer Institute Designated Massey Cancer Center grant P30 CA016059.

